# Comparative Analysis of Electrostatic Models for Ligand Docking

**DOI:** 10.1101/577643

**Authors:** Geraldo Rodrigues Sartori, Alessandro S. Nascimento

## Abstract

The precise modeling of molecular interactions remains an important goal among molecular modeling techniques. Some of the challenges in the field include the precise definition of a Hamiltonian for biomolecular systems, together with precise parameters derived from Molecular Mechanics Force Fields, for example. The problem is even more challenging when interaction energies from different species are computed, such as the interaction energy involving a ligand and a protein, given that small differences must be computed from large energies. Here we evaluated the effects of the electrostatic model for ligand binding energy evaluation in the context of ligand docking. For this purpose, a classical Coulomb potential with distance-dependent dielectrics was compared with a Poisson-Boltzmann (PB) model for electrostatic potential computation, based on DelPhi calculations. We found that, although the electrostatic energies were highly correlated for the Coulomb and PB models, the ligand pose and the enrichment of actual ligands against decoy compounds, were improved when binding energies were computed using PB as compared to the Coulomb model. We observed that the electrostatic energies computed with the Coulomb model were, on average, ten times larger than the energies computed with the PB model, suggesting a strong overestimation of the polar interactions in the Coulomb model. We also found that a slightly smoothed Lennard-Jones potential combined with the PB model resulted in a good compromise between ligand sampling and energetic scoring.

## 1 Introduction

The quantitative description of molecular interactions, at an atomic level, remains an important challenge even in current days of Petascale computing. Some of the difficulties found in this field include: (i) the energetic description of biomolecular systems; (ii) the fact that binding energies are small differences taken from large energies, resulting in large uncertainties; and (iii) the limited sampling for some calculations. Taken together, these obstacles are exactly the challenge of scoring solutions in the docking problem [1].

The second problem, due to the small differences taken from bigger numbers, can be alleviated with accurate calculations and appropriate sampling. In the context of single point calculations, such as in ligand docking, this challenge remains as an important issue and is handled in some applications with a posterior analysis of ligand candidates using molecular dynamics (MD) or Monte Carlo (MC) simulations to generate an ensemble of thermally accessible configurations of the system and binding energy calculations. In this context, the MM-GBSA or MM-PBSA approaches became very popular [2, 3, 4].

The energetic description of a biomolecular system is tackled in many docking approaches using molecular mechanics force fields [5, 6, 7], where the intermolecular interaction energies are typically computed as a sum of polar interactions, modeled as a Coulomb potential, and van der Waals interactions, modeled through a Lennard-Jones potential [8]. Additional terms can be added to model the influence of the solvent, for example [9].

Modeling polar interactions using a Coulomb potential introduces some potentially important issues. First, polarization is not considered. Although this effect might be important, a quantum description of the system would be required for appropriate treatment of the dynamics in the electron density within the active site, increasing the computational costs of the calculation. Polarization could also be taken into account by the use of polarizable force fields. However, the computational cost associated with these calculations limits their use in the context of the docking of large compound databases [10]. Second, the dielectric medium of a protein might not be exactly a constant medium, since the protein surface faces the solvent while its core might be closer to a highly hydrophobic medium. So, a representation of the electrostatic potential (and energies) as a function of a varying continuum dielectrics might be necessary, such as the treatment given by the Poisson-Boltzmann (PB) equation [11, 12, 13, 14].

Interestingly, Luty and coworkers observed that, for 20 poses of benzamidine within 8 Å of trypsin binding site, the electrostatic interaction energy computed with PB and using a simple Coulomb model assuming *ϵ* = *f*(*r*), i.e., the dielectric constant *ϵ* is a linear function of the interatomic distance *r*, showed a high correlation (*r*^2^ = 0.96) [5]. In contrary, Gilson and Honig observed that this simple distance-dependent dielectric model overestimates electrostatic interactions (also observed by Luty and coworkers) and concluded that this model does not seem to be a realistic way of treating polar interactions in biomolecular systems [15].

In late ‘80s Honig and coworkers developed the DelPhi program [16], that numerically solves the PB equation for macromolecular structures, of any shape, given atomic coordinates, atomic van der Waals parameters, and atomic charges. The calculation of electrostatic potentials within the current versions of Delphi [17, 12, 13] is fast, taking a few seconds in typical workstation computers for a small size protein. Although it might not be fast enough to be used in MD simulations, it is very competitive for docking studies, where the receptor is kept as rigid, in many strategies, and the interaction potentials can be pre-computed in grids and stored for the actual docking calculations [5, 18]. The PB calculation is under constant improvement. Recently, Li and coworkers showed that Gaussian-based smoothed dielectric function could better reproduce the assignment of PKa’s for protein residues [13]. The same approach was also applied to the ion distribution [17].

Here, we compared the results of docking enrichments and pose reproduction within the same algorithm when using Coulomb electrostatics with a distance-dependent dielectric model (i.e., *ϵ* = *r*) and using a PB electrostatic potential precomputed using DelPhi [12]. Concurrently, we evaluated the influence of Lennard-Jones soft-core potential on docking efficacy with both PB and Coulomb models. We found that the PB electrostatic model resulted in modest improvement in pose reproduction and enrichment. However, when this model was combined with a smoothed van der Waals potential, an important improvement of pose reproduction and enrichment was observed, suggesting that fine-tuning of these terms is necessary.

## 2 Methods

### 2.1 Docking Calculations

For all docking calculations reported in this work, the software `LiBELa` [19] was used. `LiBELa` (Ligand Binding Energy Landscape) uses a combination of ligand- and receptor-based strategies. For this purpose, the algorithm requires a reference ligand, that indicates the initial binding mode. The docking procedure starts with a superposition of the search ligand onto the reference ligand by using a ligand-based approach, as previously described [20]. Briefly, LiBELa describes the volume of each *i* ligand atom as a Gaussian function [20]:

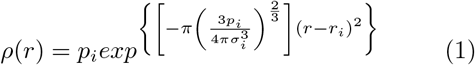

where *p_i_* is the Gaussian amplitude, defined as 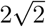, *r_i_* is atomic coordinate for atom *i* and *σ_i_* is the van der Waals radius for the same atom. Using this Gaussian-based description of shape, an overlay volume for two molecules, A and B, can be defined as [20]:

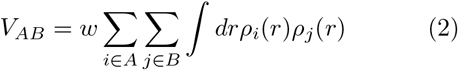

Similar terms are added to *V_AB_* to compute for the superposition of atoms with positive charge and negative charge with weights defined by *w* (here, set to 1.0 for both terms). Thus, by maximizing the overlay volume *V_AB_* in cartesian space, an initial optimized placement of the search ligand is obtained. Afterward, this initial binding mode is re-optimized to find a minimum in the binding energy using a global optimization algorithm. In this step, a typical force field-based definition of binding energy is used as the objective function:

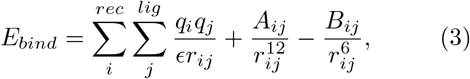

where *q* is atomic charge, *r_ij_* is the interatomic distance between atoms *i* and *j* and *A_ij_* and *B_ij_* are the Lennard-Jones parameters for the atom pair *ij*, computed by the geometric mean approximation. Here, *A_i_* = 2*δ_i_*(2*r*_0_)^12^ and *B_i_* = 2*δ_i_*(2*r*_0_)^6^, where *r*_0_ is the atomic radius and *δ* is the well depth parameter necessary for computing van der Waals interactions according to AMBER force field. Both parameters are taken from AMBER FF14SB force field [21]. A final similarity index can be computed using a Hodgkin’s similarity index [22] defined as:

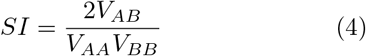

To speed up the calculations, the receptor interaction potential is precomputed and stored in grids. In this point, `LiBELa` can compute a typical Coulomb electrostatic potential:

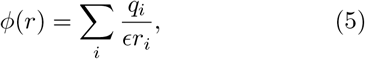

where the dielectric constant was set to the interatomic distance *r*, i.e, *ϵ* = *r_ij_* [5]. Alternatively, LiBELa can parse a DelPhi electrostatic map with the electrostatic potential *ϕ_DelPhi_* instead and compute binding energies using this stronger electrostatic model. For the calculations shown in this work, DelPhi 8.4 was used [12, 13]. Typically, a computation box of 30 × 30 × 30 Å with a spacing of 0.4 Å (gsize 75 and scale 2.5 Å, in Delphi parameters), with interior dielectrics of 2.06 and exterior dielectrics of 78.5, and salt concentration set to 145 mM. The same grid spacing was used in calculation employing the Coulomb model.

We also tested the effect of a smoothed Lennard-Jones potential by applying the same strategy as suggested by Verkhivker and coworkers [23]. Here, the binding energy is evaluated as [9]:

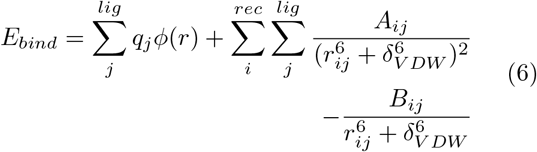

The smoothing term *δ_V DW_* was systematically varied in the interval 0.5 to 2.5 Å with a step of 0.5 Å to evaluate the effect of the Lennard-Joned soft-core potential in pose reproduction and enrichment when combined with a Coulomb electrostatic potential (*ϕ_Coulomb_*) or a PB electrostatic potential (*ϕ_DelPhi_*).

### 2.2 Docking Pose Reproduction

#### 2.2.1 Self-docking test

For docking pose reproduction, we used three data sets. The dataset SB2012 [24] includes 1,043 crystal structures of protein-ligand complexes, distributed as SYBYL MOL2 files. Here and all over this text, the ‘receptor’ is defined as a protein where an organic small molecular, the ‘ligand’, binds. In these files, the atomic charges are already defined using AMBER forcefield for receptor and AM1-BCC [25, 26] for ligands. The dataset files were used as provided, with no further optimizations or modifications of atomic coordinates. Here, a docking calculation was set using each ligand-receptor pair, using the own ligand as the reference ligand in LiBELa.

#### 2.2.2 Cross-docking test

For a cross-docking experiment, the Astex dataset was used [27]. In this dataset, 58 structures with analogous complexes are provided. From this dataset, 54 targets were used together with 860 ligands in total. The targets (receptors) were prepared using DockPrep tool as available in UCSF Chimera [28] using AMBER FF14SB atomic charges. For the ligands, AM1-BCC atomic charges were attributed using ANTECHAMBER [29] and SYBYL atom types were assigned using the same tool. In this experiment, each ligand was docked on different (non-native) crystal structures of its own target. Afterward, the root mean square deviation (RMSD) was computed using the native ligand structure as a reference.

### 2.3 Enrichment Tests

In order to evaluate the ability to enrich actual ligands against decoys, i.e., compounds with similar physicochemical properties but not expected to bind to a given target, the DUD38 subset of DUD-E database, which contains 38 targets from the original DUD dataset [30], but rebuild with the same protocol as used in DUD-Enhanced (DUD-E) [31]. This subset includes the PDB files for the receptors and over 630,000 compounds, among binders and decoys, with an average decoy-to-ligand ratio of 33. The compounds were used as provided (as SYBYL MOL2 files) with atomic charges defined following the default ZINC protocol [32, 33]. The receptor files were prepared using the DockPrep tool available in UCSF Chimera [28]. In this tool, atomic charges are attributed to receptor atoms following AMBER FF14SB parameters. Finally, the prepared receptor is saved as a SYBYL MOL2 file type.

The target-specific ligands and decoys were docked to each target using LiBELa default parameters and using either a Coulomb electrostatic model or a pre-computed Delphi electrostatic potential. The Delphi calculations were carried out in two steps. In the first step, a calculation is set where the protein represents 50% of the calculation box. In a second step, a focused calculation was carried out using a grid of 0.4 Å for a 30 × 30 × 30 Å calculation box centered in the center of mass of the reference ligand. The energies computed after docking calculations were used to rank the docked molecules and ROC curves were computed with locally developed python scripts. The enrichment was quantified using the Adjusted LogAUC metric [34]. This metric is similar to the well-known AUC but is computed for a semi-logarithmic plot of the ROC curve spanning three decades in the horizontal axis. The computed area is then corrected to remove the area expected for a random enrichment (14.5%).

## 3 Results

The calculations of the electrostatic potentials with DelPhi are very fast, typically taking less than five seconds in an Intel Xeon E5645 (2.40GHz) processor running in a single thread. This is much faster than the calculation of the interaction potential grids in LiBELa, which took about 5.4 minutes averaging over the 38 targets of the DUD38 dataset. The computational efficiency of the electrostatic calculations with DelPhi makes it tempting to use this more robust model in docking calculations. However, what is the actual role of PB-based calculation on protein-ligand interactions in the context of ligand docking? In order to address this question, we set up a comparative analysis of ligand binding poses and ligand enrichments using the Coulomb electrostatic model or PB as the electrostatic model to assess the effect of the model in sampling and enrichment, respectively.

### 3.1 Effect on Sampling

The simplest experiment one can think of to evaluate the sampling effect on ligand docking is to assess the ability of a model to reproduce ligand poses from crystal structures. In this context, we compared the root mean square deviations observed for LiBELa using either Coulomb or PB as the electrostatic model.

After the redocking of 1,029 ligands on their respective receptors, the RMSD for all atoms, including hydrogen atoms, was computed in comparison with the original (experimental) structures. Averaging over the entire dataset an RMSD of 1.215 Å was observed for the Coulomb model, while for the PB model an average RMSD of 1.129 Å was achieved. The median RMSD for these models were 0.535 and 0.598 Å with a standard deviation slightly increased for Coulomb as compared to PB (1.689, compared to 1.419 Å). For both models the fraction of targets with RMSD values found below the typical cutoff value of 3.0 Å was close to 90%, as indicated in Table 1.

**Table 1:** Summary of the self-docking experiment using the SB2012 dataset. N=1,030. Statistics for RMSDs computed after redocking of ligands are shown in comparison with experimental poses. HA_RMSD, HA_RMSDh and HA_RMSDm refer respectively to the average standard RMSD computed for heavy-atoms only, the minimum-distance heavy-atom RMSD used in Autodock Vina [35] and Hungarian (symmetry-corrected) heavy-atom RMSD [36], as computed by DOCK6 [37].

| Electrostatic Model | Coulomb | PB |
| --- | --- | --- |
| Smoothing Parameter $\delta_{VDW} = 0.0$ Å | | |
| Average RMSD (Å) | 1.215 | 1.129 |
| Median RMSD (Å) | 0.535 | 0.598 |
| Standard Deviation (Å) | 1.689 | 1.419 |
| < 3.0 Å | 90% | 92% |
| Average HA_RMSD (Å) | 1.124 | 1.35 |
| Average HA_RMSDh (Å) | 1.035 | 0.948 |
| Average HA_RMSDm (Å) | 0.534 | 0.494 |
| Average Hodgkin’s SI | 0.858 | 0.871 |
| Smoothing Parameter $\delta_{VDW} = 0.5$ Å | | |
| Average RMSD (Å) | 1.287 | 1.079 |
| Median RMSD (Å) | 0.535 | 0.596 |
| Standard Deviation (Å) | 1.812 | 1.312 |
| < 3.0 Å | 88% | 93% |
| Average HA_RMSD (Å) | 1.192 | 0.987 |
| Average HA_RMSDh (Å) | 1.104 | 0.901 |
| Average HA_RMSDm (Å) | 0.538 | 0.467 |
| Average Hodgkin’s SI | 0.852 | 0.875 |
| Smoothing Parameter $\delta_{VDW} = 2.0$ Å | | |
| Average RMSD (Å) | 5.643 | 1.185 |
| Median RMSD (Å) | 5.566 | 0.611 |
| Standard Deviation (Å) | 2.304 | 1.525 |
| < 3.0 Å | 15% | 91% |
| Average HA_RMSD (Å) | 5.356 | 1.096 |
| Average HA_RMSDh (Å) | 4.881 | 1.007 |
| Average HA_RMSDm (Å) | 2.263 | 0.527 |
| Average Hodgkin’s SI | 0.287 | 0.858 |

When a smoothed Lennard-Jones potential was combined with the electrostatic models under evaluation in this work, we found very interesting differences. For a small smoothing parameter *δ* = 0.5Å, the differences between the electrostatic models are small, similarly to what is observed in the AMBER Lennard-Jones model. However, as *δ* becomes larger, the differences between the Coulomb model and the PB model become more evident. When *δ* is set to 2.0 Å, the average RMSD found for the Coulomb model was 5.643 Å (median 5.566 Å), while the average RMSD for the PB model was 1.185 Å, with median in 0.611 Å. So, it appears that the combination of the PB model with a soft-core VDW potential still leads to good results in pose reproduction while the Coulomb model rapidly seems to dominate the binding energy resulting in meaningless ligand poses.

Another interesting observation comes from the comparison between the polar term in the interaction energies. An analysis for 1,029 protein-ligand complexes reveals a good correlation between the electrostatic interaction energies computed using a Coulomb model with distance-dependent dielectrics, i.e., *δ* = *r_ij_*, and electrostatic interaction energies computed using the Poisson-Boltzmann model. As shown in Figure 1, there is a good correlation between the computed energy terms (*r* = 0.7 *for N* = 1, 029), as also observed previously by Luty and coworkers [5]. Additionally, one can observe that the electrostatic interaction energies computed by the Coulomb model are about 10 times more favorable, on average than those computed using the PB model, indicating a typical overestimation of the interaction energies in this model. In the context of ligand docking, this overestimation may result in binding modes that are biased towards a few polar contacts that are too favorable as compared to the overall fitting of the ligand and receptor binding pockets.

**Figure 1:**
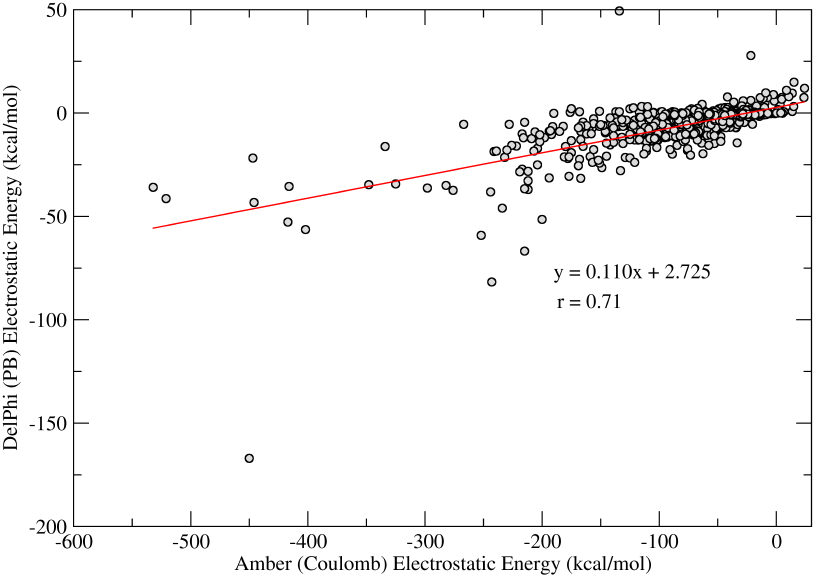
Correlation of the electrostatic interaction energies computed with Coulomb model (horizontal axis) and PB (vertical axis). The line shows a linear regression of the obtained data (*N* = 1,029) and the regression coefficients are shown in the figure.

A more stringent test is to assess the ability of the model to reproduce experimentally determined poses in a cross-docking experiment, i.e., in a receptor structure different from the one used in the actual docking calculation. In brief, it involves docking ligand *L_A_* in receptor structure *R_B_* and comparing the docking pose to the pose observed when *L_A_* was crystallized bound to receptor *R_A_*. For this task, the Astex non-diverse dataset was used.

The employed dataset includes 603 diverse (non-native) complexes. The results obtained are summarized in Table 2. Again, when the typical AMBER Lennard-Jones potential is used, a slight improvement in the binding poses is observed, with average RMSD going from 4.48 Å, for the Coulomb model, to 3.95 Å in the PB model (median values of 3.27 and 2.94 Å, respectively). When a smoothed Lennard-Jones potential is used, on the other hand, the differences between both electrostatic models increases. For a smoothing parameter *δ* set to 0.5 Å, the average RMSD decreases from 4.65 in the Coulomb model to 3.90 in the PB model (median values 3.52 and 3.06 Å). And when *δ* is set to 2.0 Å, the average RMSD decreases from 7.60 Å to 3.71 Å, with median values decreasing from 7.52 to 2.54 Å (Table 2).

**Table 2:** Summary of the cross-docking experiment using the Astex dataset. N=603. Statistics for RMSDs computed after redocking of ligands in a non-native receptor structure are shown in comparison with experimental poses.

| Electrostatic Model | Coulomb | PB |
| --- | --- | --- |
| Smoothing Parameter $\delta_{VDW} = 0.0$ Å | | |
| Average RMSD (Å) | 4.477 | 3.951 |
| Median RMSD (Å) | 3.269 | 2.938 |
| Standard Deviation (Å) | 4.408 | 4.091 |
| Average HA_RMSD (Å) | 4.194 | 3.698 |
| Average HA_RMSDh (Å) | 3.768 | 3.272 |
| Average HA_RMSDm (Å) | 4.075 | 3.801 |
| Average Hodgkin's SI | 0.533 | 0.562 |
| Smoothing Parameter $\delta_{VDW} = 0.5$ Å | | |
| Average RMSD (Å) | 4.653 | 3.900 |
| Median RMSD (Å) | 3.523 | 3.064 |
| Standard Deviation (Å) | 4.537 | 4.014 |
| Average HA_RMSD (Å) | 4.362 | 3.635 |
| Average HA_RMSDh (Å) | 3.851 | 3.270 |
| Average HA_RMSDm (Å) | 4.074 | 3.790 |
| Average Hodgkin's SI | 0.521 | 0.562 |
| Smoothing Parameter $\delta_{VDW} = 2.0$ Å | | |
| Average RMSD (Å) | 7.603 | 3.712 |
| Median RMSD (Å) | 7.519 | 2.542 |
| Standard Deviation (Å) | 3.758 | 4.053 |
| Average HA_RMSD (Å) | 7.369 | 3.457 |
| Average HA_RMSDh (Å) | 6.721 | 3.077 |
| Average HA_RMSDm (Å) | 5.699 | 3.688 |
| Average Hodgkin's SI | 0.206 | 0.579 |

Taken together, the results shown here indicate that the PB model for electrostatic computation result in better pose reproduction in the scenario of a typical AMBER FF binding energy calculation and, more significantly, in the scenario of a soft docking, i.e., when the Lennard-Jones potential is smoothed.Given the results obtained, we moved for the evaluation of the changes in the enrichment of actual binders against decoy compounds.

### 3.2 Ligand Enrichment

In order to assess the ability of the electrostatic models to recover actual ligands against decoy compounds, we choose the DUD38 dataset. In this dataset, 38 targets are given with a set of binder compounds and a set of decoy compounds. In this context, a decoy is defined as a compound that has similar physicochemical properties to the binders but is not expected to bind to the receptor. After docking all the binders and decoys, the compounds are ranked by their binding energy and a receiver-operating characteristic (ROC) curve is plotted. Finally, the enrichment is computed using Adjusted LogAUC metric, as previously proposed [34]. Briefly, this metric describes the area under the curve of the ROC plot with the x-axis in the logarithm scale and spanning three decades. The area computed is corrected by subtracting the area expected for a random enrichment.

The results obtained are summarized in Table 3 and shown in the complete version in the Supplementary Material. From the data shown here, we note that, for the usual Lennard-Jones model used in AMBER force field, i.e., *δ_V DW_* = 0.0, the electrostatic models performed almost similarly in terms of enrichment, with an average enrichment of 4.50 or 4.91 for Coulomb or PB respectively, with a slight improvement of the enrichment with the PB model. Using the smoothed VDW potential with *δ_V DW_* = 0.5Å, similar enrichments are observed but with an improvement in the median enrichment for the PB model. Here, the average enrichments were 5.01 and 4.99 with median enrichments of 3.93 and 5.40 for Coulomb and PB models, respectively. Finally increasing the smoothing constant to *δ_V DW_* = 2.0Å, a maximum in the average/median logAUC is observed for the PB model (5.36 and 6.09 for average and median, respectively), while a marked decrease in the enrichment for the Coulomb model is observed.

**Table 3:** Summary of the enrichment experiment using the DUD38 dataset. The values are reported as Adjusted LogAUC and also as AUC, in the parenthesis.

| Electrostatic Model | Coulomb | PB |
| --- | --- | --- |
| Smoothing Parameter $\delta_{VDW} = 0.0\text{\AA}$ | | |
| Average | 4.50 (56.2%) | 4.91 (57.3%) |
| Median | 3.24 (54.9) | 4.70 (56.0%) |
| Standard Deviation | 6.87 (10.4%) | 4.42 (8.3%) |
| Smoothing Parameter $\delta_{VDW} = 0.5\text{\AA}$ | | |
| Average | 5.01 (57.1%) | 4.99 (57.5%) |
| Median | 3.93 (56.4%) | 5.40 (57.2%) |
| Standard Deviation | 6.81 (10.5%) | 4.03 (7.7%) |
| Smoothing Parameter $\delta_{VDW} = 2.0\text{\AA}$ | | |
| Average | 1.80 (53.4%) | 5.36 (58.7%) |
| Median | 0.52 (52.2%) | 6.09 (58.6%) |
| Standard Deviation | 5.14 (8.5%) | 3.87 (7.4%) |
| DOCK 6.7 Grid Score |  |  |
| Average | 1.3 (42.5%) |  |
| Median | -1.2 (43.3%) |  |
| Standard Deviation | 9.7 (17.4%) |  |

For the sake of comparison, the same docking calculations using the DUD38 were set up using the Grid Score model of DOCK 6.7 [37]. The average and median logAUC observed for this model was 1.3 and −1.2, respectively (Table 3). Since logAUC corrects for the expected random enrichment, this metric can achieve negative results if results are worse than random. It is important to add that the Grid Score here used a 6-12 Lennard-Jones potential with a Coulomb electrostatic model that uses a distance-depend dielectric function (*ϵ* = *r_ij_*), similar to the model used in LiBELa.

## 4 Discussion

The enrichment data shown in Table 3 for the DUD38 dataset strongly suggests that the continuum electrostatic model can lead to important improvements in the ability to recover actual binders and separate them from decoy compounds. On the other hand, as we already noted from the data shown in Figure 1, the Coulomb interaction electrostatic energies are, on average, 10 times more favorable than interaction electrostatic energies computed with PB. Then, it makes sense that the balance between the electrostatic and van der Waals terms should be also fine-tuned. We assessed this balance by introducing a smoothed Lennard-Jones term to model the van der Waals interactions. A good balance seems to be achieved when the smoothing constant *δ_V DW_* was set to 2.0 Å. With this calculation setup, a maximum in the enrichment is observed, without compromising the docking poses, according to the results of ligand enrichment with DUD38, self-docking with the SB2012 dataset (Table 1) and cross-docking with the Astex dataset (Table 2). A complete comparison of the effect of the smoothing parameter *δV DW* is shown in the Supplementary Material, where the ligand pose and ligand enrichment can be compared as a function of the smoothing parameter.

A second effect of the electrostatic treatment given to the docking calculations can be observed in the distribution of the net charges of the top-scored molecules in docking calculations. The analysis of the charge distribution for the target ace, shown as an example in Figure 2, reveals that among the top-scored molecules when the Coulomb model was used, almost half of them have net charges −2 or −3 e, indicating a favoring of the non-specific electrostatic interactions to the total docking score. On the other hand, the PB model favors neutral molecules or molecules with net charge −1 e. No molecule with net charge −2 or −3 is observed among the top-scored molecules, suggesting a much more specific scoring of the biomolecular interactions. As a piece of evidence of the correctness of the PB model, an inspection of the distribution of net charges among the actual binders in the DUD dataset for this target shows that 66% of the binders have net charge 0, 30% have charge −1 and 4% have charge −2, indicating that the PB model more closely reflects the molecular interactions observed in experimental conditions.

**Figure 2:**
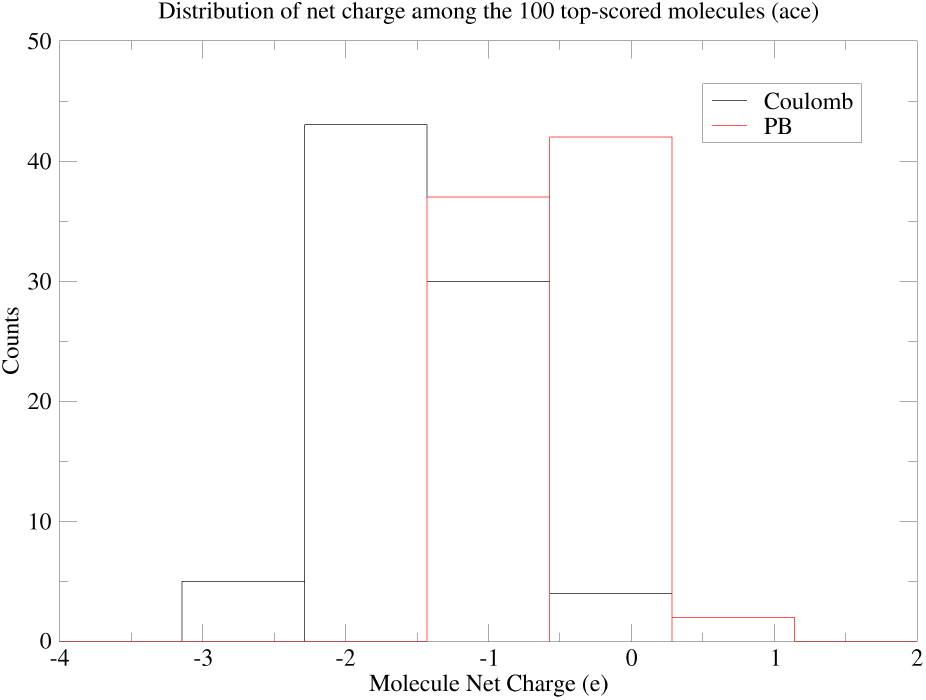
Distribution of the net charge for the 100 top-scored molecules in docking calculations of the DUD38 target ace. The calculations were done without soft-core potential, i.e., with smoothing parameter *δ_V DW_* set to 0 Å.

A recent development in the PB calculations introduced a Gaussian-based approach ‘to deliver a smooth dielectric function for the entire space domain’ [13]. The authors showed that the Gaussian-based function resulted in better assignments of PKa’s and also dielectric values for protein interior and protein-water interface in agreement with previous works [13]. Interestingly, a comparison of the ligand enrichment obtained after docking calculations using the Non-Gaussian dielectric model and the Gaussian dielectric model showed a decrease of about 15% in the enrichment of actual ligands against decoys in the DUD38 dataset. This decrease is observed for the AMBER Lennard-Jones model as well as for the smoothed Lennard-Jones model. This still preliminary observation highlights that even the robust PB model can still be improved in the context of ligand docking to result in even more reliable calculations and predictions of protein-ligand interactions.

In conclusion, here we evaluated the effect of scoring docking calculations with a Coulomb model or with a Poisson-Boltzmann model for electrostatic energies. We found that equally good docking poses are observed in self-docking calculations. However, the more stringent test of cross-docking calculations indicated an improvement of the docking pose when the PB model was used as compared to the Coulomb model. Finally, the enrichment of actual binders as compared to decoy compounds was improved when the PB model was used and balanced with smoothed van der Waals interactions. Together the results shown here suggest that better models are computationally viable, in terms of time and efficiency, and the effects of the improvements in the model can dramatically affect the outcome of docking calculations, with great potential for the screening of drug candidates.

## 5 Acknowledgments

The authors thank the `DelPhi` developers for making their tool freely available to the scientific community. We also thank Heloisa Muniz and Victor Nogueira for the discussions. We thank the financial support provided by Fundação de Amparo à Pesquisa do Estado de São Paulo (FAPESP) through grants 2017/18173-0, 2015/26722-8 and 2015/13684-0 and by Conselho Nacional de Desenvolvimento Científico e Tecnológico (CNPq), through grants 303165/2018-9 and 406936/2017-0.

