## Supplementary Data for "Comparative Analysis of Electrostatic Models for Ligand Docking"

Supplementary Material

### 1.1 Supplementary Table

**Supplementary Table 1**. Summary of the self-docking, cross-docking and enrichment experiments. The self- and cross-docking results are shown as the average RMSDs over 1,030 SB2012 targets (self-docking) and 603 Astex non-native targets (cross-docking). The enrichment results are shown as the average Adjusted LogAUC over the 38 targets of the DUD38 subset of DUD-E.

| **δ_VDW_** | **Electrostatic Model** | **Self-Docking** | **Cross-Docking** | **Enrichment** |
| --- | --- | --- | --- | --- |
|  | | **RMSD (Å)** | | **logAUC** |
| 0.5 | PB | 1.079 | 3.900 | 4.99 |
| 2.0 | PB | 1.185 | 3.712 | 5.36 |
| 0.0 | PB | 1.129 | 3.951 | 4.91 |
| 0.0 | Coulomb | 1.215 | 4.477 | 4.50 |
| 0.5 | Coulomb | 1.287 | 4.653 | 5.01 |
| 2.0 | Coulomb | 5.643 | 7.603 | 1.80 |

### 1.2 Supplementary Figures

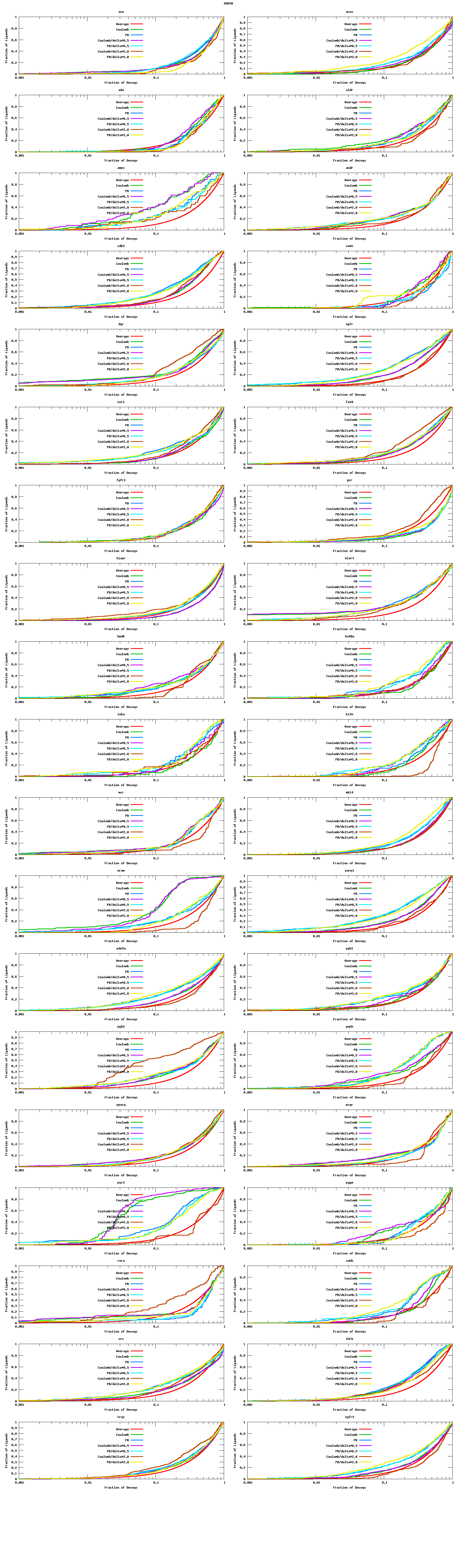

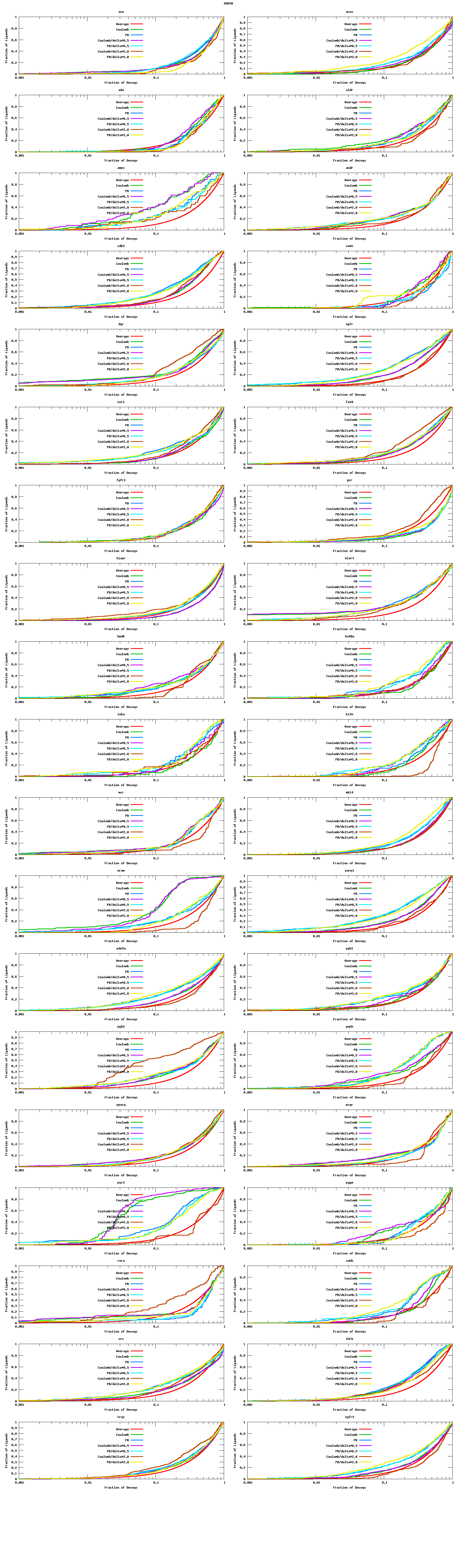

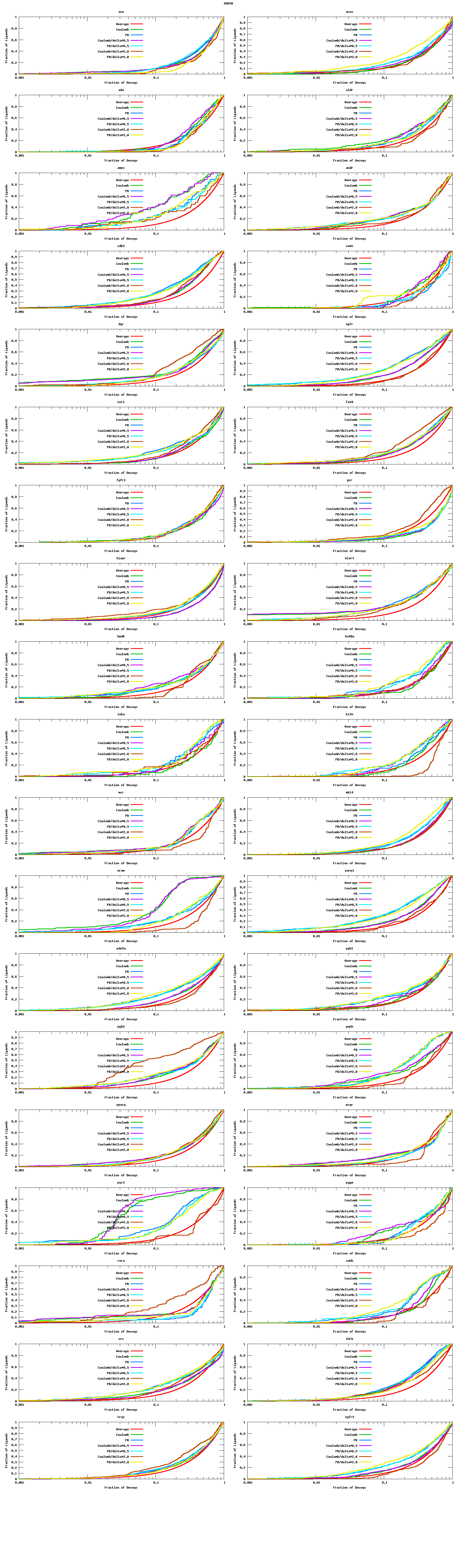

**Supplementary Figure 1.** Enrichment plots with adjusted logAUC (area under the curve) of known ligands against decoys for the 38 targets of DUDE38 database and for different docking condition.

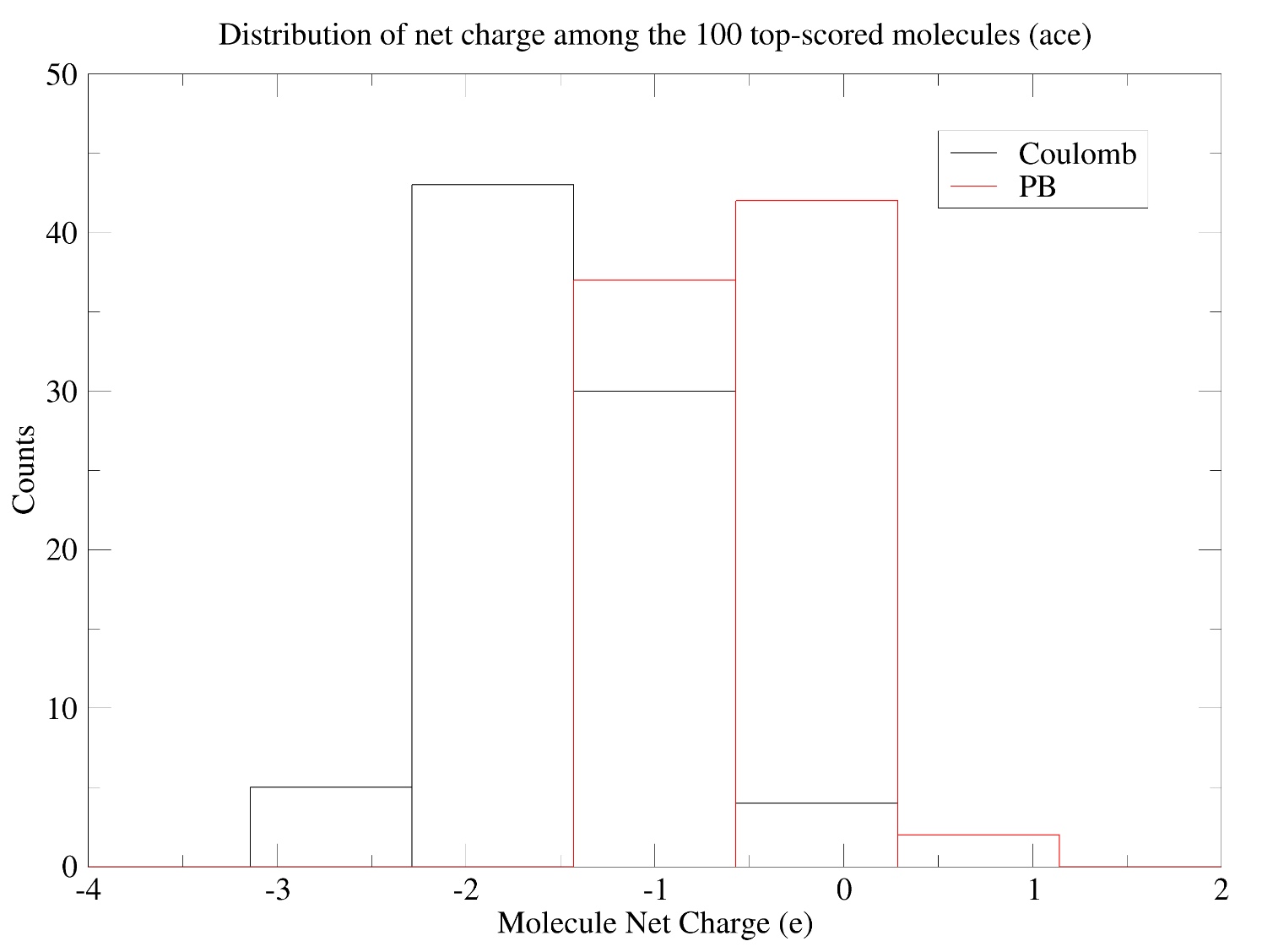

**Supplementary Figure 2.** Distribution of the net charge for the 100 top-scored molecules in docking calculations of the DUD38 target ace. The calculations were done without soft-core potential, i.e., with smoothing parameter δ_VDW_ set to 0 Å.
